# Establishment of the trans-resveratrol-induced mouse pulmonary fibrosis model and its comparison with the bleomycin-induced model

**DOI:** 10.1101/812883

**Authors:** Chen Yiwen, Zong Chenzhong, Sheng Chunrui, Chang Hongsheng, Wang Shuyan, Zhao Hongzhao, Liu Shanshan, Zhang qiaohui, Dong Ruijuan, Ge Dongyu, Yu Xue, Li Lina

## Abstract

Pulmonary fibrosis (PF) is a progressive, fatal disease, and its pathogenic mechanism has not yet been identified. The bleomycin (BLM)-induced animal model has several shortcomings that cannot be overcome. We accidentally found that the continuous oral administration of trans-resveratrol (TR) at levels above the prescribed dose can induce PF within a certain period of time, and this model can be successfully replicated in mice, as indicated by the hydroxyproline (HYP) test and pathological analysis. The TR model requires no anesthesia or surgery and thus causes less damage to mice than the BLM model. The progress of fibrosis in the TR model was slow but progressed steadily compared with the BLM model, and the TR model was more comparable to human PF. Although the pathogenesis is not understood, the TR-induced PF model indicated that there may be a close relationship between PF and tumors and antitumor drugs, which requires further exploration.

## Introduction

Pulmonary fibrosis (PF) is a progressive, fatal disease, and its pathogenic mechanism has not yet been determined. Therefore, the generation of an animal model that can accurately replicate this disease is of great significance for research on the mechanisms and clinical treatment of this disease. An ideal animal model should resemble human disease as closely as possible, be easy to perform, be widely available, be less expensive than current models, and most importantly, be highly reproducible and consistent. Trans-resveratrol (TR), which is derived from resveratrol^1^, is a polyphenolic compound that has multiple effects. TR has a better antiradical efficacy than resveratrol^2^ and can reduce inflammation, regulate cell signaling, induce apoptosis by upregulating or downregulating protein expression, cause growth arrest via the cell cycle, and resist apoptosis and oxidation^3^. Currently, the trans form is frequently used as a health-care product to protect against the incidence of cardiovascular pathologies^4^. During clinical diagnosis and treatment, we accidentally found that the continuous oral administration of TR drugs at levels above the prescribed dose can induce PF in patients within a certain period of time.

Thus far, we have successfully modelled this effect in mice with a TR drug 4 times and observed that the model differs from the classical bleomycin (BLM) model, but its pathogenesis is still not fully understood. Therefore, we compared this model with the BLM model and tried to explore whether this animal model was more analogous to the pathological process in human PF to further elucidate the pathogenesis of this disease and to develop more tools to develop clinical treatments.

## Materials and Methods

### 1. Materials

#### 1.1. Reagents

TR (Vitacost, US) was dissolved at a concentration of 4.29% in 0.5% carboxymethyl cellulose sodium (Sigma-Aldrich, US). To obtain a dose of 57.14 mg/kg body weight (BW), 0.008 ml/g BW of 4.29% TR solution was administered by gavage. BLM hydrochloride for injection (Haizheng-Huirui, China) was dissolved at a concentration of 2.5 mg/ml with sterile 0.9% NaCl (isotonic saline) for intratracheal administration. The injected BLM solution volume was 2 μl/g BW to obtain a dosage of 5 mg/kg BW.

#### 1.2. Animals

Four to five-week-old male Institute of Cancer Research (ICR) mice weighing 18–22 g were purchased from SPF China (Beijing, China) and were kept under specific pathogen-free conditions, 12-h light/dark cycles, and constant temperature with food and water available ad libitum at the Beijing University of Chinese Medicine animal facility. Animals were allowed to acclimate in the facilities for at least 3 days before receiving any treatments. The procedures for the care and use of animals were approved by the Experimental Animal Ethics Committee of Beijing University of Traditional Chinese Medicine, and all applicable institutional and governmental regulations concerning the ethical use of animals were followed.

### 2. Methods

#### 2.1. Models of BLM-induced and TR-induced PF

##### 2.1.1. Establishment of the BLM-induced mouse model

More than 84 mice were included in this group. All of these mice were anesthetized with intraperitoneal injections of pentobarbital sodium salt (1% concentration; Sigma, US) at 40 mg/kg BW. Following anesthesia, a midline cut of the neck skin was made, and the trachea was exposed by blunt dissection. The needle of a 1-ml syringe was inserted into the trachea, and BLM was dissolved in 50 μl sterile saline and was injected into the lungs at a dose of 5 mg/kg BW. The mice were rotated immediately after the injection to ensure that BLM was evenly distributed in the lungs, and then the neck skin incision was sutured.

##### 2.1.2. Establishment of the TR-induced model in mice

More than 84 mice were included in this group. A 0.008 ml/g BW solution of 4.29% TR was administered by gavage. Mice received TR by gavage once a day from day 1 to day 28. For time course experiments, lung samples were collected on days 7, 14, 28, 35, 42 and 56 for further analysis.

The process of the experiment was as follows.

- Week 0. Mice were randomly divided into 3 groups. In consideration of the experimental mortality, each group contained at least 84 mice but not more than 100 mice. The TR model, BLM model, and control groups were established for comparison. The TR model was administered drug for 28 consecutive days, and the BLM model was treated once. The control group was treated with a normal feeding method.
- Week 1. Fourteen mice were randomly selected from each group and were sacrificed, 7 for bronchoalveolar lavage fluid (BALF) analysis and 7 for lung tissue analysis.
- Week 2. Fourteen mice were randomly selected from each group and were sacrificed, 7 for BALF analysis and 7 for lung tissue analysis.
- Week 4. Fourteen mice were randomly selected from each group and were sacrificed, 7 for BALF analysis and 7 for lung tissue analysis.
- Week 5. Fourteen mice were randomly selected from each group and were sacrificed, 7 for BALF analysis and 7 for lung tissue analysis.
- Week 6. Fourteen mice were randomly selected from each group and were sacrificed, 7 for BALF analysis and 7 for lung tissue analysis.
- Week 8. Fourteen mice were randomly selected from each group and were sacrificed, 7 for BALF analysis and 7 for lung tissue analysis.

#### 2.2. Extraction of experimental materials

Mice were randomly chosen from each group and were anesthetized with intraperitoneal injections of pentobarbital sodium salt. After visible anesthetization, mice were immobilized on the mouse plate in the supine position. The abdominal cavity was opened along the median line. The abdominal aorta was severed to bleed before killing the mice. Then, the thoracic cavity was quickly opened, the lungs were exposed, the left and right lung lobes were removed, and the residual blood was absorbed with filter paper. The wet weights of the whole lungs were measured. Then, the lungs were placed in the corresponding cryopreservation tubes and were stored in the freezer at −80°C.

#### 2.3. Lung histology

The middle lobe of the right lung was removed en bloc and was placed in 20 ml of 10% formaldehyde for 48 hours. Then, the lungs were subsequently dehydrated with ethanol and xylene and were embedded in paraffin. Four-micron-thick sections were cut and stained with hematoxylin and eosin (H&E) and Masson dyes. Five microscopic fields of the right middle lung were randomly chosen and microscopically photographed with a 20-fold magnification in all experimental mice by a technician who was blinded to the results (three photographs per mouse). The authors chose 108 photographs according to quality criteria (sharply photographed and >95% of the photograph had to contain lung tissue) and ensuring that the frequency of each grade of fibrosis was similar to avoid creating a bias for certain fibrotic grades. The degrees of alveolitis and fibrosis were determined as described by Ashcroft^5^ and Hübner^6^.

#### 2.4. Hydroxyproline (HYP) measurement

First, a standard curve of HYP was generated. Eight centrifuge tubes were used to make different concentrations (0, 3.90625, 7.8125, 15.625, 31.25, 62.5, 125 and 250 μg/ml) of an HYP standard solution. Then, 200 μl of 0.1 mol/l citric acid buffer solution and 0.05 mol/l chloramine T solution were added to these centrifuge. After fully mixing the contents of the centrifuge tubes, the tubes were placed in an incubator with a constant temperature and were cultured at 37°C for 15 minutes. After the incubation, 200 μl of 3.15 mol/l perchloric acid solution was added to each centrifuge tube, and the contents were mixed and were then incubated at 37°C for 5 minutes. Then, 200 μl of 10% 4-(dimethylamino)benzaldehyde (PDAB) solution (dissolved in anhydrous ethanol) was added to each centrifuge tube, and the contents were thoroughly mixed. Then, the tubes were incubated in an 85°C water bath for 3 minutes. When the incubation was completed, the centrifuge tubes were cooled with ice water, and 200 μl of the contents was removed from each tube and added to an enzyme label plate. The plate was placed in the enzyme label instrument, and the absorbance was determined at 560 nm. The standard curve was drawn according to these results.

The right upper lobe was weighed. The tissues were dried at 70°C until their weight no longer decreased. Each tissue homogenate was prepared with 2 mol/l sodium hydroxide solution (10%). The homogenate was moved to a centrifuge tube, a hole was drilled in the cap and the tube was wrapped in tin foil, then the tissue was digested under 0.1 kPa pressure for 20 minutes at 120°C. After cooling to room temperature, the homogenate was centrifuged for 10 minutes at a speed of 12,000 revolutions per minute. Fifty microliters of supernatant from every sample was taken and placed into a new centrifuge tube. The next steps were the same as that of the standard curve. The level of HYP was calculated according to the standard curve. The reagents were used within 7 days after preparation.

#### 2.5. The collection of BALF

Mice were anesthetized, and their tracheae were exposed. A plastic cannula was inserted into the trachea. Then, 0.5 ml of cold phosphate buffered saline (PBS) was injected into the lungs through the cannula, flushed back and forth, and then as much was as possible. This step was repeated three times. Approximately 1 ml of BALF was collected and immediately stored at 4°C. The cell pellet was further processed to classify and count the cells with an automated hematological analyzer Sysmex XS-800i. Briefly, the white blood cells in alveolar lavage fluid were counted. The proportions of neutrophils, lymphocytes, monocytes, eosinophils and basophils were evaluated.

#### 2.6. Test of the effect on lung oxidant/antioxidant biomarkers

The samples used for these tests were from the right lower lobe of the lung in mice. First, every sample was processed to generate a homogenate supernatant with a concentration of 10%. Each sample was weighed and placed in a glass homogenizer and was then mixed with cold control saline (9 times the volume of the lung). The mixture was ground on ice until it generated tissue homogenate. Then, the homogenate was transferred into a centrifuge tube and was centrifuged for 10 minutes at a speed of 12,000 revolutions per minute at 4°C. After centrifugation, the supernatant was used for the following experiment.

The optimal homogenate concentration to detect the three indicators was determined according to the results of preliminary experiments. The concentrations required for malondialdehyde (MDA), SOD and GSH-PX detection were 10%, 0. 1% and 1.75%, respectively. Therefore, the supernatant of the 10% homogenate was diluted to these three concentrations with cold control saline before the experiment. The detection reagent was purchased from Nanjing Jiancheng Institute of Bioengineering. The protocols for the detection these three indicators were performed strictly in accordance with the kit instructions.

#### 2.7. The expression of relevant cell factors in the lung tissue

Several major cell factors were measured, such as Tumor Necrosis Factor alpha (TNF-α), Interferon-gamma (IFN-γ), Interleukin-4 (IL-4), IL-12p70 and IL-13. The same samples were used as in the as antioxidant/oxidant analyses. The experimental procedure was carried out in strict accordance with the ELISA kit instructions. The kits were purchased from Invitrogen.

#### 2.8. Statistical analysis

A total of 5 to 7 values from each group were included in the statistical analysis. Data are presented as the mean ± the standard deviation (SD). Statistical analyses and graphing were performed using GraphPad Prism 7 software (GraphPad Software Inc., San Diego, CA, USA). The following statistical tests were conducted: two-way analysis of variance (ANOVA) followed by Tukey-Kramer’s multiple comparisons and the Kruskal - Wallis test followed by Dunn’s test for parametric and nonparametric measures, respectively. Values of p < 0.05 were considered statistically significant.

## Results

1. Lung coefficient, weight and mortality of mice (Figure 1).
2. Lung histology, i.e., H&E and Masson staining (Figure 2).
3. The expression of HYP in the lung tissue (Figure 3).
4. The cell classification and counts in BALF (Figure 4).
5. The effect on lung oxidant/antioxidant biomarkers (Figure 5).
6. The expression of relevant cell factors in the lung tissue (Figure 6).

**Figure 1.**
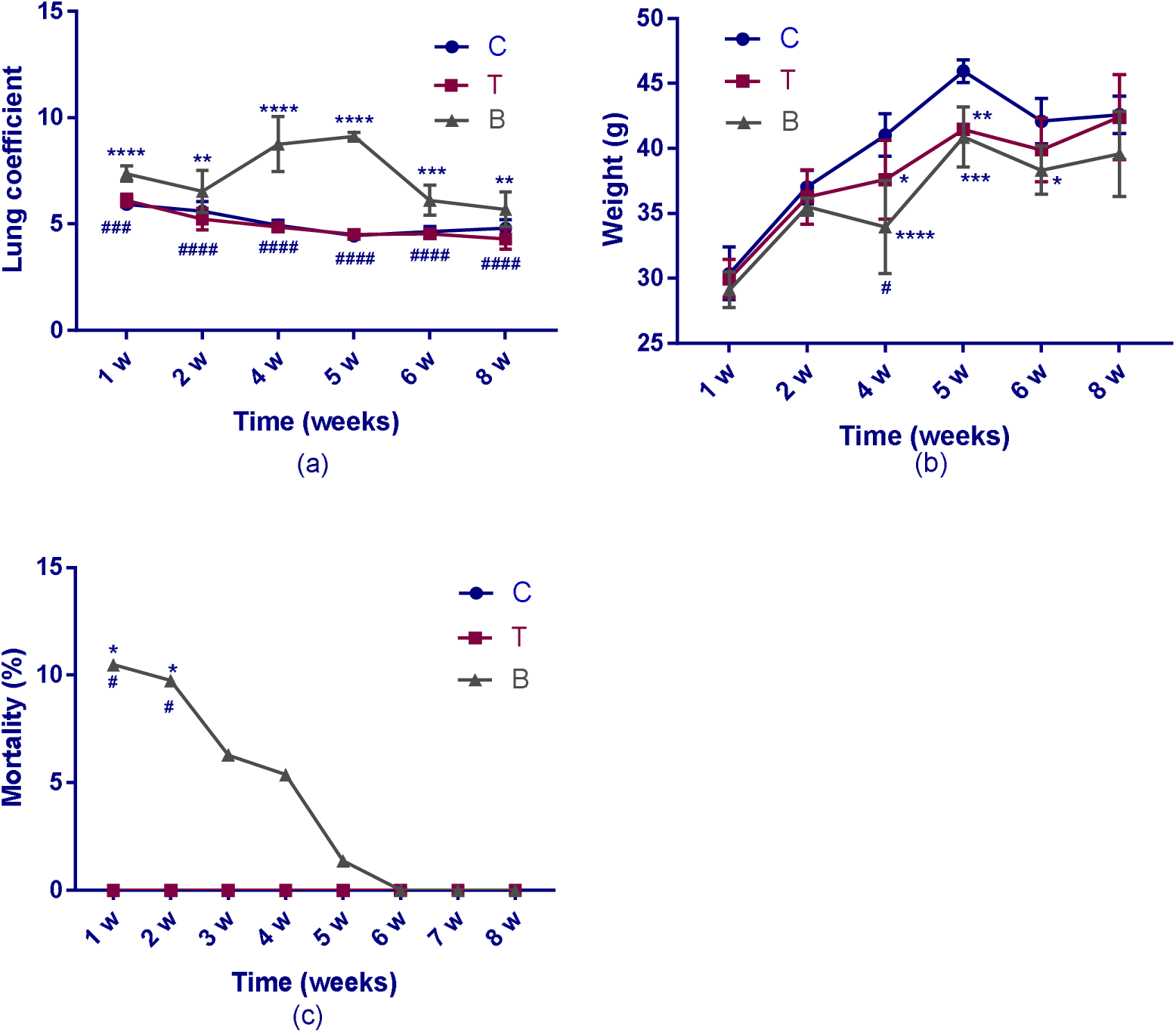
(a) Lung coefficient at 1, 2, 4, 5, 6 and 8 weeks. No differences were observed for the lung coefficient in the control group and the TR group, and the trends in the changes in the two groups were consistent. However, significant differences were observed between the control group and the BLM group, as well as between the TR group and the BLM group. A more obvious difference was observed in the 4th and 5th weeks. (b) Weight of the mice at 1, 2, 4, 5, 6 and 8 weeks. Statistical analysis showed that there was a marked difference among the three groups. The weight of the mice in the control group was the highest, followed by the weight of the mice in the TR group, and the weight of the mice in the BLM group was the lowest. The first decrease in the BLM group appeared in the 4th week. All the groups exhibited weight decreases by the 6th week, but the weight increased in the 8th week. (c) Mortality of the mice at 1, 2, 3, 4, 5, 6, 7 and 8 weeks. The graph shows that the mortality of the BLM group was distinctly higher than that of the other groups, especially in the first two weeks. Later, the difference between the three groups gradually decreased, and the mortality of the three groups eventually became identical. C represents the control group, T represents the TR group, and B represents the BLM group. **** *p*<0.0001, *** *p*<0.0002, ** p<0.0021, and * *p*<0.0332 compared with the control group; #### *p*<0.0001, ### *p*<0.0002, ## p<0.0021, and # p<0.0332 compared to the BLM group. For the statistical analysis, two-way ANOVA followed by Tukey’s multiple comparison test was used.

**Figure 2A.**
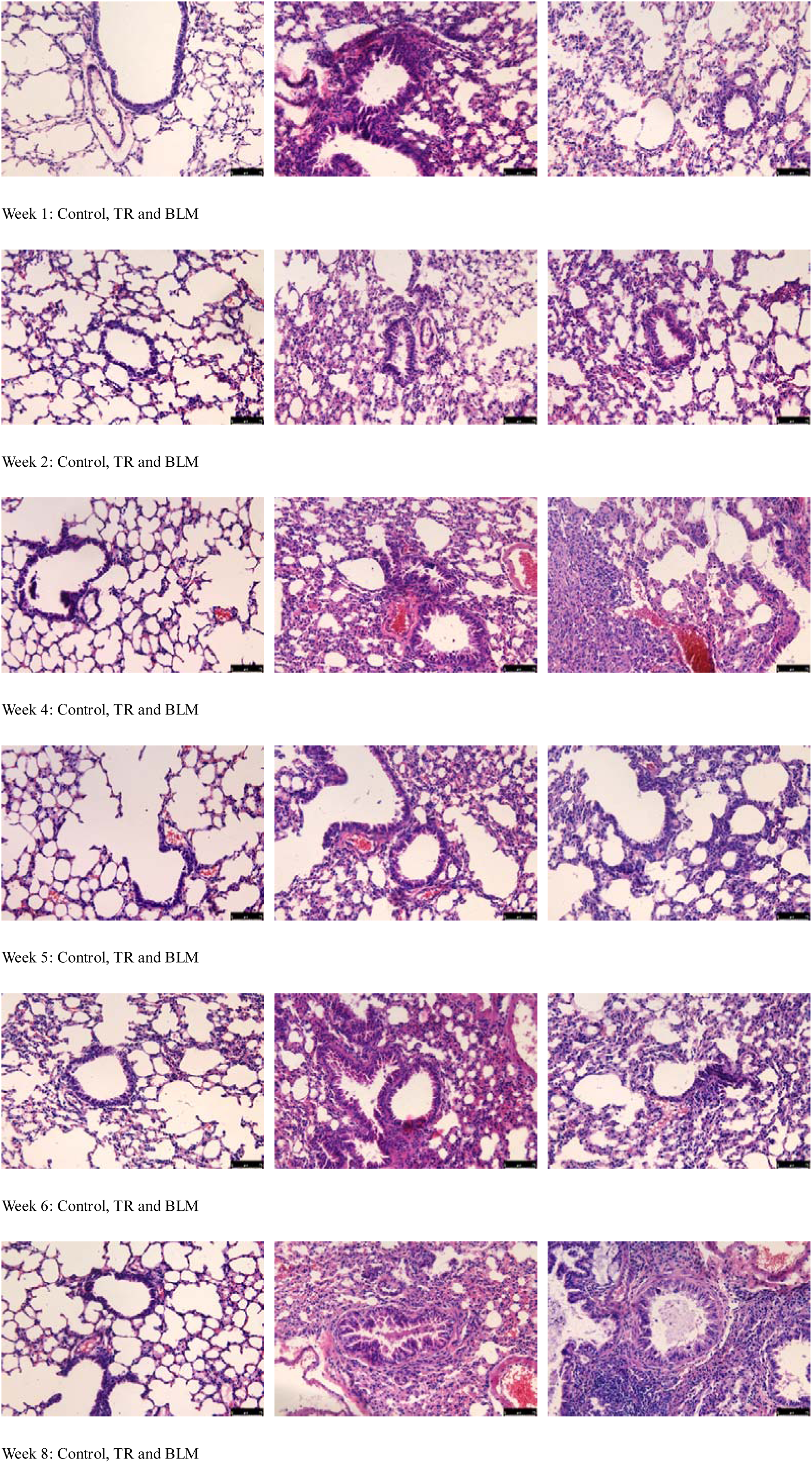
Lung histology at 1, 2, 4, 5, 6 and 8 weeks. H&E staining (200×). The results showed that the control group had intact lung architecture with no evidence of inflammation, edema, hemorrhage, emphysema or fibrosis at all timepoints. The TR group showed moderate hemorrhages, emphysema, areas of increased thickening of the alveolar septa, leukocytic infiltration in the alveolar walls, and fibroplasia. However, the BLM group showed more severe pathological changes overall.

**Figure 2B.**
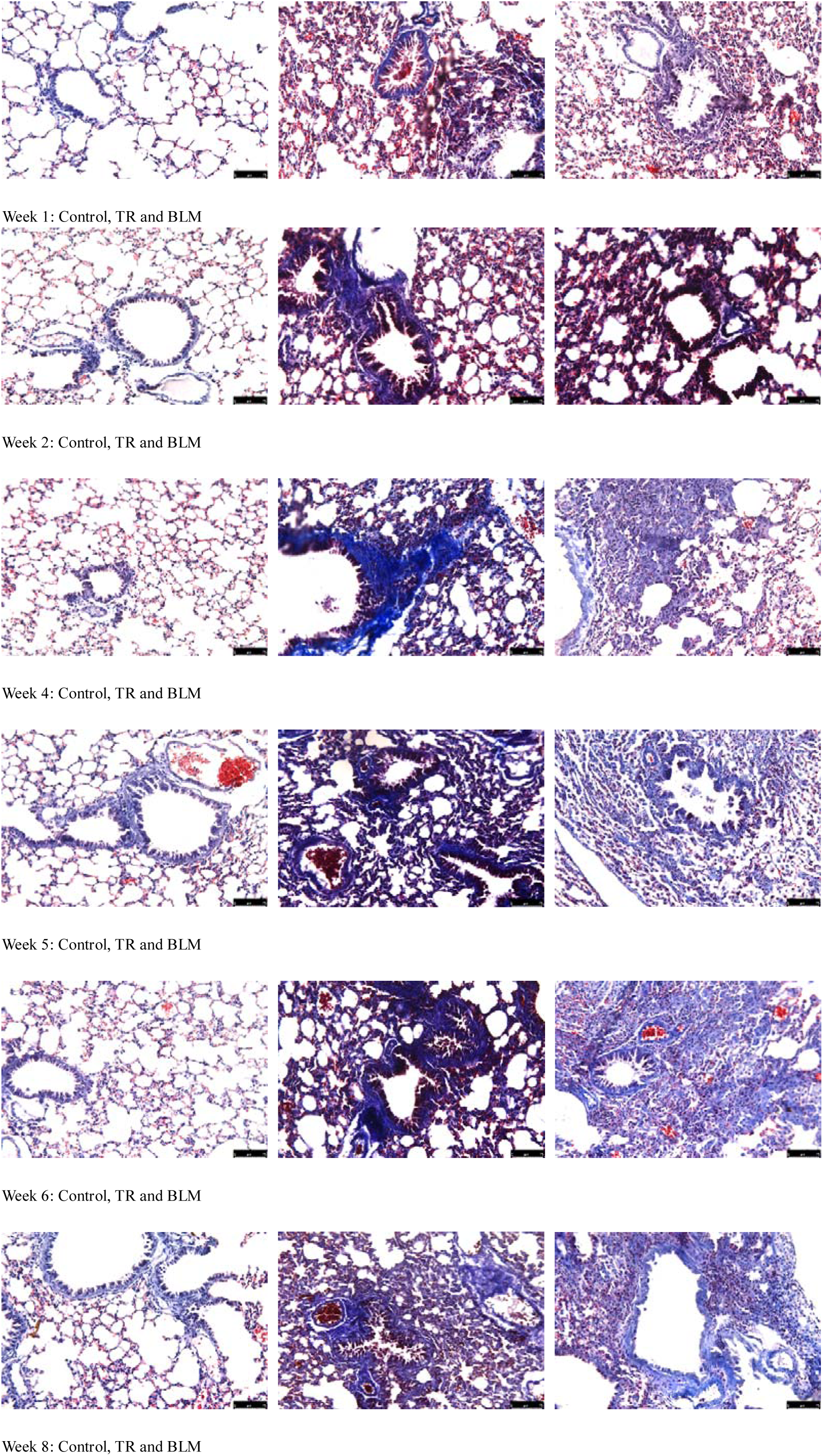
Lung histology at 1, 2, 4, 5, 6 and 8 weeks. Masson staining (200×). The results showed the control group had no blue staining, indicating that there was no collagen deposition in interstitium as well as in the alveolar septae. The TR group showed bluish staining, indicating that there was collagen deposition in the interstitial space and the peribronchial area, while the BLM group showed similar but more severe pathological changes.

**Figure 3.**
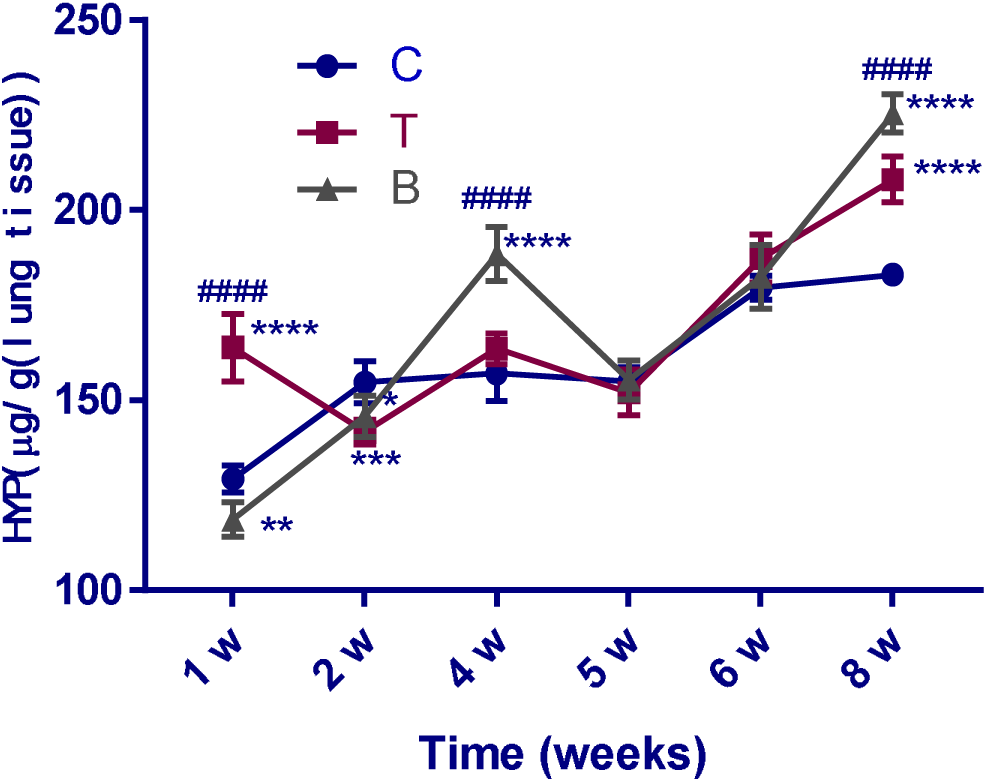
HYP content in the lungs of mice at 1, 2, 4, 5, 6 and 8 weeks. The results showed that there was a notable difference among the three groups (p<0.0001). For the TR and BLM groups, significant differences appeared in the 1st, 4th and 8th weeks. For the control and the other two groups, the most prominent difference occurred in the 1st, 2nd, 4th and 8th weeks. Interestingly, there was no clear distinction among these groups in the 5th and 6th weeks. C represents the control group, T represents the TR group, and B represents the BLM group. **** *p*<0.0001, *** *p*<0.0002, ** p<0.0021, and * p<0.0332 compared with the control group; #### *p*<0.0001, ### *p*<0.0002, ## p<0.0021, and # p<0.0332 compared to the BLM group. For the statistical analysis, two-way ANOVA followed by Tukey’s multiple comparison test was used.

**Figure 4.**
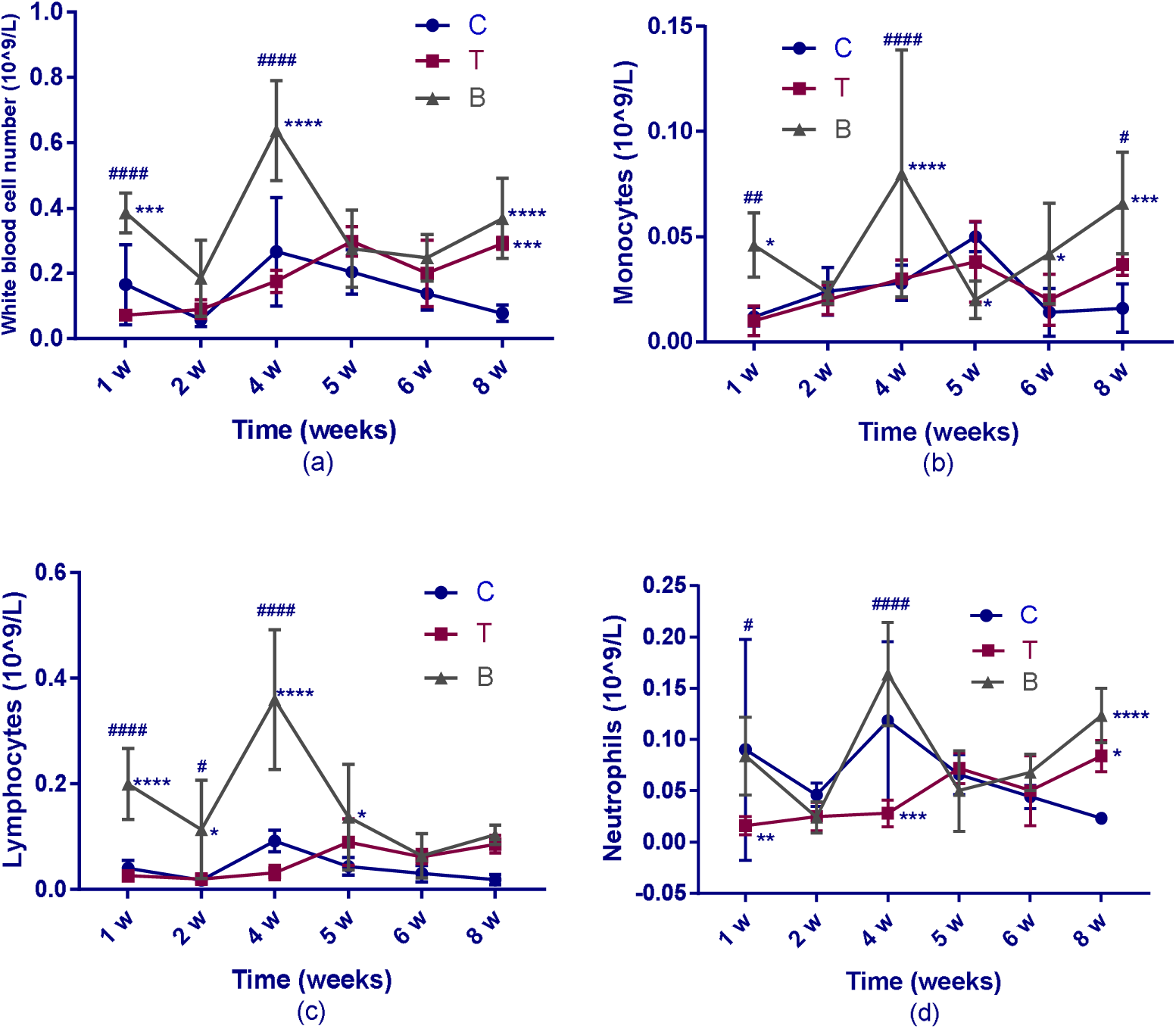
(a) The number of white blood cells in the BALF of mice at 1, 2, 4, 5, 6 and 8 weeks. The results showed a remarkable difference among the control group, TR group and BLM group at every time point. Compared with the control group, the TR group showed a significant difference at the 8th week, while the BLM group showed a significant difference at the 1st, 4th, and 8th weeks. When classifying the relevant white blood cells, the results showed that the monocytes (b), lymphocytes (c) and neutrophils (d) did not change, and almost the same trend was observed in the control group. In contrast, the BLM group greatly fluctuated, as did the white blood cells in this group. It is worth noting that the neutrophils in the control group fluctuated more than those in the TR group, and a similar trend was observed in the BLM group. C represents the control group, T represents the TR group, and B represents the BLM group. **** *p*<0.0001, *** *p*<0.0002, ** p<0.0021, and * p<0.0332 compared with the control group; #### *p*<0.0001, ### *p*<0.0002, ## p<0.0021, and # p<0.0332 compared to the BLM group. For the statistical analysis, two-way ANOVA followed by Tukey’s multiple comparison test was used.

**Figure 5.**
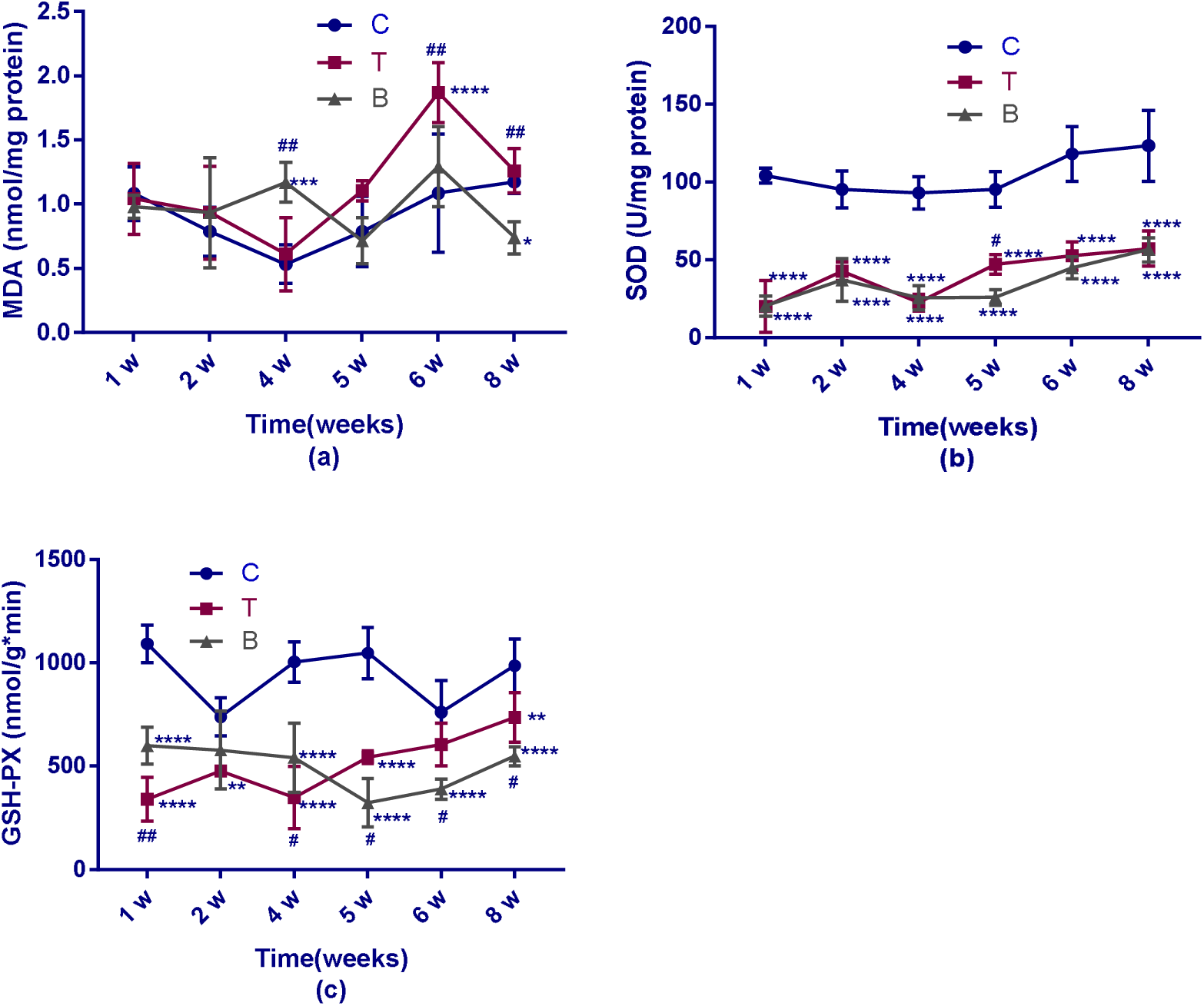
The MDA (a), SOD (b) and GSH-PX (c) levels in the lungs of mice at 1, 2, 4, 5, 6 and 8 weeks. The MDA (a) levels were not notably different in the three groups in the early phase, but a large difference appeared in the later phase, especially at the 4th week and later. The MDA content of the BLM group was higher than that of the other groups at the 4th week, while the MDA level was the highest in the TR group starting at the fifth week. It is worth noting that the SOD level (b) of both the TR group and the BLM group was significantly lower than that of the control group. Meanwhile, the levels in the TR and BLM groups were not significantly different. The GSH-PX content (c) was different in all three groups, both in terms of the trends in the changes and the specific values. The results of the control group remained higher than those in the other two groups during all 8 weeks. The two model groups, however, had changing trends. Specifically, the TR group surpassed the BLM group and was then surpassed by the BLM group in the former and latter 4 weeks, respectively. C represents the control group, T represents the TR group, and B represents the BLM group. **** *p*<0.0001, *** *p*<0.0002, ** p<0.0021, and * p<0.0332 compared with the control group; #### p<0.0001, ### p<0.0002, ## p<0.0021, and # p<0.0332 compared to the BLM group. for the statistical analysis, two-way ANOVA followed by Tukey’s multiple comparison test was used.

**Figure 6A.**
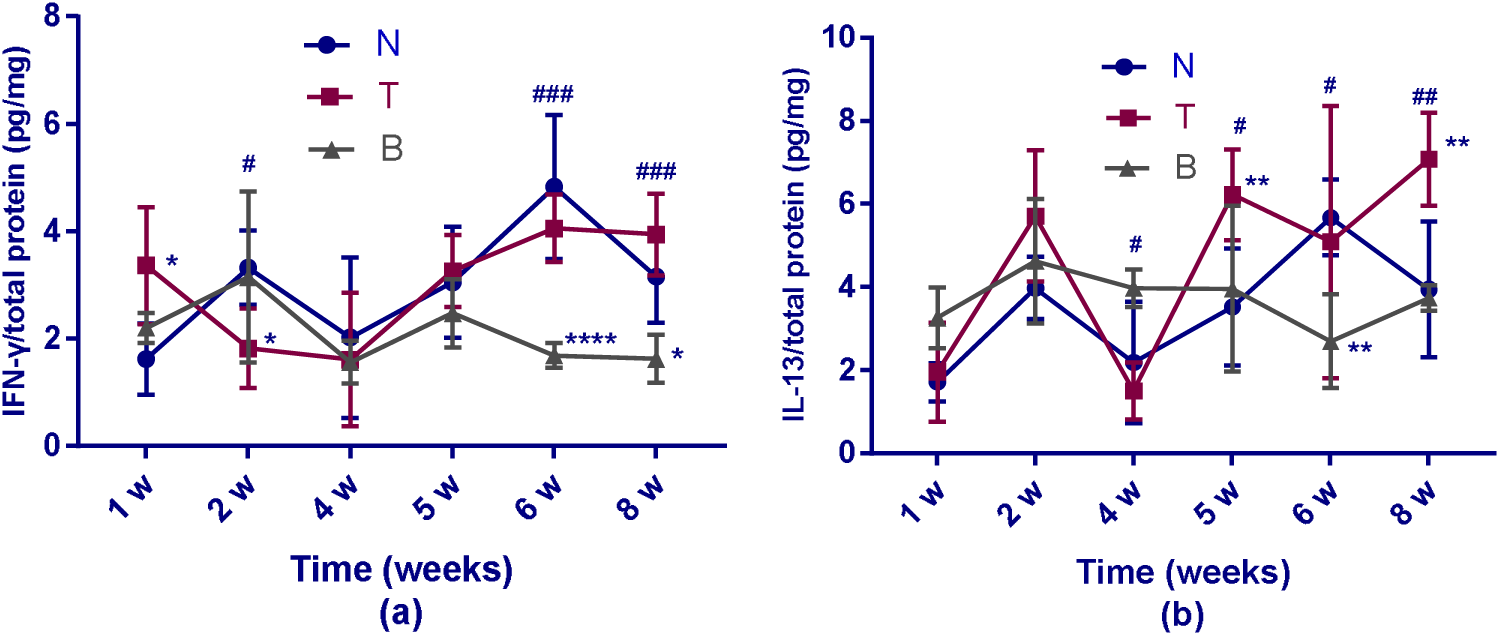
Expression of IFN-γ (a) and IL-13 (b) in the lungs of different groups at 1, 2, 4, 5, 6 and 8 weeks. The trend in IFN-γ (a) showed more obvious differences among the three groups (*p*<0.0002). In the 1st week, the expression of the TR group was highest, while it became the lowest the next week. In the latter two weeks, the expression in the BLM group remained the lowest. For the two model groups, the difference in the expression appeared only in the 2nd, 6th and 8th weeks. Specifically, the expression in the TR group was lower in the former phase but was higher than that in the BLM group in the latter phase. Nevertheless, the difference between the control and the two model groups was not large. The trend in IL-13 (b) showed a relatively significant distinction among the three groups (*p*<0.0021), but the overall trend was complex and irregular. The expression in the TR group significantly surpassed that in the control group only in the 5th and 8th weeks, while in the 6th week, the expression in the BLM group was lower than that in the control group. When comparing the expression in the two model groups, the statistical analysis showed that slight distinctions occurred from the 4th to 6th week, and then the distinction became greater in the 8th week. During the phases mentioned above, the expression in the TR group rebounded and even peaked, while the expression in the BLM group fluctuated once but was stable overall. C represents the control group, T represents the TR group, and B represents the BLM group. **** *p*<0.0001, *** *p*<0.0002, ** p<0.0021, and * p<0.0332 compared with the control group; #### *p*<0.0001, ### *p*<0.0002, ## p<0.0021, and # p<0.0332 compared to the BLM group. For statistical analysis, two-way ANOVA followed by Tukey’s multiple comparison test was used.

**Figure 6B.**
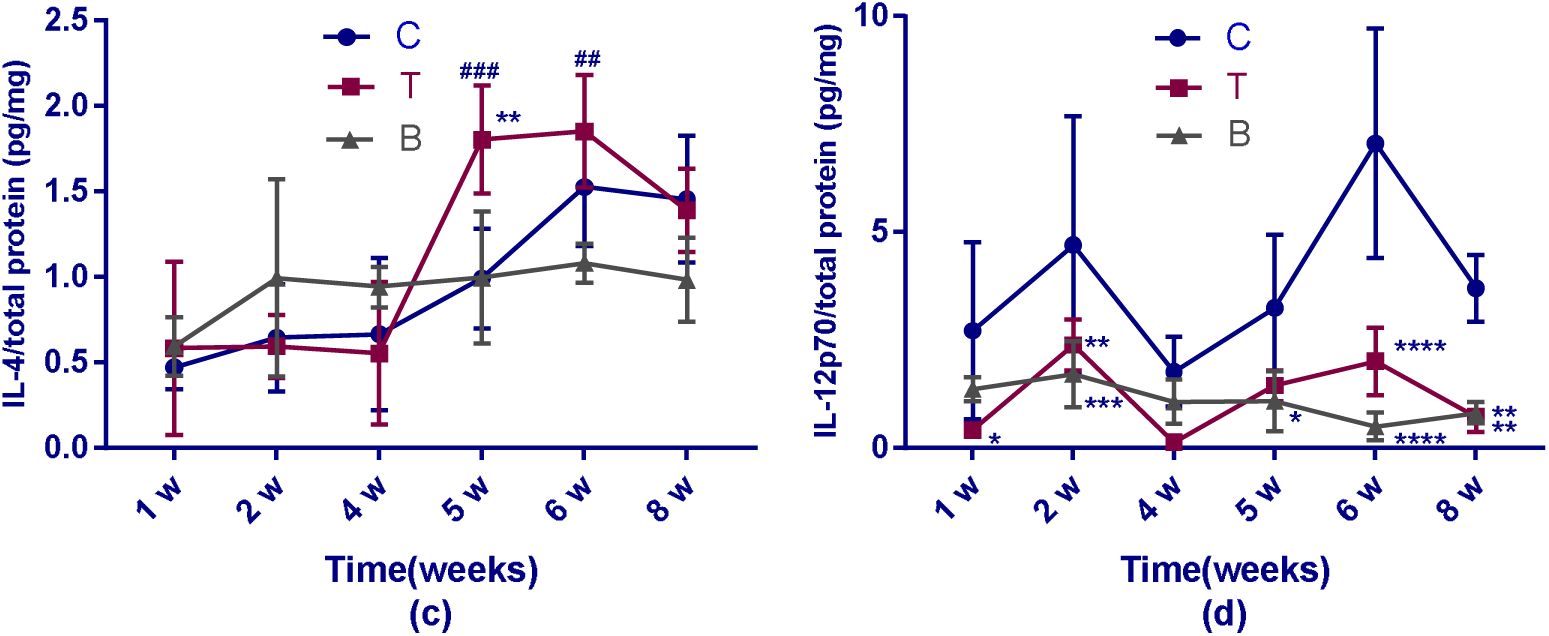
Expression of IL-4 (c) and IL-12p70 (d) in the lungs of different groups at 1, 2, 4, 5, 6 and 8 weeks. The trend in IL-4 (c) showed no notable difference among the three groups. Of the two model groups, the TR group significantly surpassed the control group only in the 5th week. For the two model groups, the TR group was higher than the BLM group in the 5th and 6th weeks. The trend in IL-12p70 (d), however, showed a clear distinction between the control and the two groups; in other words, the expression of the two groups remained lower than that of the control group (*p*<0.0001). Of all the weeks tested, the most significant difference was in the 6th week, and the next most significant difference was in the 2nd week, followed by the 8th week. Meanwhile, it is worth noting that the expressions of the two model groups was not different during the whole period. C represents the control group, T represents the TR group, and B represents the BLM group. **** *p*<0.0001, *** *p*<0.0002, ** p<0.0021, and * p<0.0332 compared with the control group; #### *p*<0.0001, ### *p*<0.0002, ## p<0.0021, and # p<0.0332 compared to the BLM group. For the statistical analysis, two-way ANOVA followed by Tukey’s multiple comparison test was used.

**Figure 6C.**
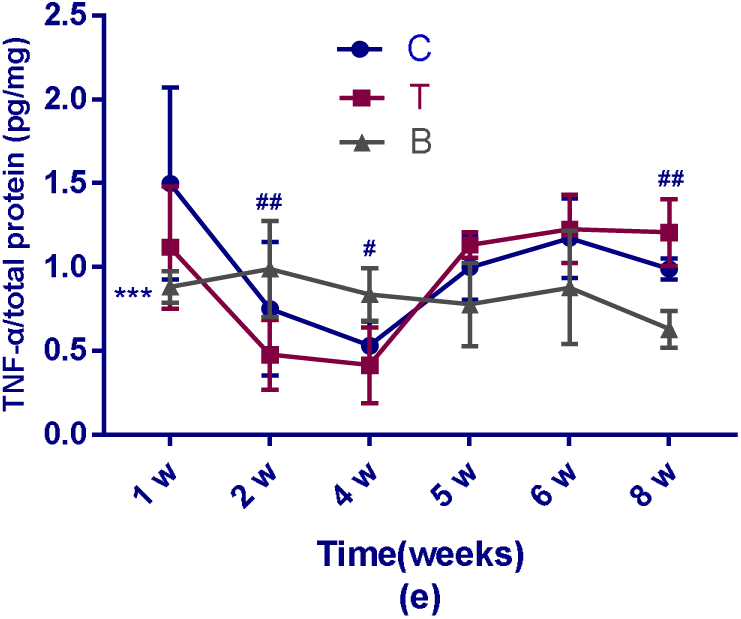
Expression of TNF-α (e) in the lungs of different groups at 1, 2, 4, 5, 6 and 8 weeks. The trend in TNF-α (e) was not notably different among the three groups, which was especially clear when comparing the TR and control groups, while the trend in the BLM group was the opposite to the other two groups. In the 1st week, the expression in the TR group was significantly lower than that in the other two groups. In the 2nd and 4th weeks, the difference in the expression between the TR and BLM groups became clear, and the level in the BLM group remained higher. However, the level of the two model groups decreased in the 8th week. C represents the control group, T represents the TR group, and B represents the BLM group. **** *p*<0.0001, \*\*\**p*<0.0002, ** p<0.0021, and * p<0.0332 compared with the control group; #### *p*<0.0001, ### *p*<0.0002, ## p<0.0021, and # p<0.0332 compared to the BLM group. For the statistical analysis, two-way ANOVA followed by Tukey’s multiple comparison test was used.

## Discussion

### 1. Analyses of the results of the TR model indicators and its comparison with the BLM model

As the PF development process in the BLM model has been relatively well studied, we compared the BLM model with the TR model to examine the pathological changes in our new model as much as possible. It has been experimentally proven that animals in the BLM model usually develop fibrotic lesions around the 2nd to the 6th week^7^**Error! Bookmark not defined.**, yet some researchers found that the fibrosis can be maintained at 6 months^8^.

#### 1.1. Lung histology

Starting in the first week of modeling, the TR group showed lesions consistent with alveolitis and PF, which included the thickening of the alveolar septum, a massive deposition of collagen, and scattered inflammatory secretions in the lumen of some of the lung tissue. In the 2nd week, the collagen deposition in the TR group was generally lower than that in the previous week, and the alveolar structure was mostly intact. The BLM model began to progress from acute inflammation to fibrosis, pulmonary tissue hemorrhage began to decrease, the alveolar septum began to thicken, and collagen deposition began to occur. By the 4th week, both models were developing fibrosis, and the collagen deposition significantly increased compared with that in the previous week, but the degrees of collagen deposition and alveolar septal damage in the TR model were lower than those in the BLM model. At the 5th week, the fibrosis process of the two models slowed down again. In the 6th week, the models began to progress toward PF, and the degree of the lesions in the TR model was slightly more severe than that in the BLM model. At the end of the 8th week, the fibrosis levels of the two models reached their highest levels in the experiment, but level in the TR model became lower than that in the BLM model again.

#### 1.2. The HYP content

The HYP level is usually considered one of the gold standards for evaluating PF^9^, because the continuous accumulation of collagen is the main pathological manifestation of fibrosis^10^. In our experiment, there was a notable difference in the HYP content among the three groups. As the graph in Figure 3 shows, the HYP contents in the BLM model and the TR model tended to be similar in the later stages of the experiment; at the end of the experiment, the levels of these two groups were significantly higher than those of the control group as well as of the BLM model at the 4th week. When we evaluated this result in association with the results of pathological section staining, we concluded that both TR and BLM generated a PF model in this strain of mice, and the difference between the two groups was in the degree of fibrosis and the process of disease development.

#### 1.3. The results of sorting and counting the white blood cells in BALF

In addition to the above two gold-standard indicators, we also tested other indicators that are commonly used in animal models of PF. By sorting and counting the white blood cells in BALF, we found that the number of white cells in the BLM model was always higher than that in the TR model and the control group, indicating that there was more obvious inflammation in the BLM model at this stage, which was basically consistent with the reported trend. In the TR model, the number of white blood cells fluctuated only slightly during the whole experiment, which may indicate that there was no obvious inflammatory phase in the pathological process of this PF model. By counting the monocytes and lymphocytes in the white blood cells, it was shown that the BLM model had the largest fluctuation, while the TR model always slightly fluctuated around the level of the control group. This may indicate that the progress of the TR model was connected with “monocyte to macrophage differentiation”^11^. The neutrophil counts showed that there was a significant fluctuation in the TR model throughout the experiment. This may indicate that the TR model causes neutrophil recruitment^12^.

#### 1.4. The oxide and antioxidant contents in the lung tissue

We also tested the levels of several oxides and antioxidants in the lung tissue of mice in both models. MDA is a product of lipid peroxidation^13^. SOD is an antioxidant metalloenzyme present in living organisms; GSH-PX is an important peroxide-degrading enzyme that is widely present in the body, and both of these enzymes are a part of the antioxidant defense system^14^. The results showed that the MDA content of the TR model increased sharply starting in the 4th week and then fell returned to the level of the control group at the 6th week. The BLM model and the control group had only small fluctuations. In contrast to the MDA level, the SOD and GSH-PX content were significantly lower in both the TR and BLM models than in the control group, indicating that both the TR and BLM drugs produced a large number of free radicals. Therefore, a large amount of SOD and GSH-PX were consumed, which indicates these drugs induced oxidative stress in mice^15^. According to previous studies, when a high levels of free radicals are produced in the body, the MDA content tends to increase^16^. However, in this experiment, we found that when the SOD content of the two models decreased, the MDA content at the corresponding time point did not synchronously increase, but both peaked and fell after the fourth week. Based on this observation, we suspect that in the first four weeks, the body could compensate, and the measured MDA levels were not high. When the generated MDA exceeded the compensatory capacity of the body, the measurement suddenly increased.

#### 1.5. The expression of cytokines related to the PF model

In addition, we also examined the expression levels of several cytokines that are important in the PF model. The type 2 cytokine response as characterized by the cytokines IL-4 and IL-13, and type 1 cytokine responses are characterized by IL-12 and IFN-γ, in immunity, inflammation, and fibrosis, and both responses exhibit remarkable crossregulation or inhibit the opposite pathway^17^. Previous studies have shown that the levels of TNF-α^18^, IFN-γ^19^, IL-12^20^, IL-4 and IL-13^21^ are generally upregulated in the lung tissue of mice with BLM-induced PF. However, it is worth considering whether our experimental results were not consistent with the literature reports. We need to further explore what factors caused these conditions.

### 2. The significance of the new TR model

#### 2.1. The defects of the BLM model

Based on our knowledge, it is widely recognized that BLM can be used to induce replicable animal models of PF that are similar to PF in humans^22^. BLM is a chemotherapeutic drug that is used to treat a variety of neoplastic diseases. The intratracheal administration of BLM can cause central fibrotic changes in bronchioles, acute interstitial and alveolar inflammation, macrophage activation and the upregulation of TNF-α, granulocyte macrophage colony-stimulating factor and some interleukins^23^. However, in terms of the similarity of the evolution of the disease, the BLM model shows rapid progress and reversible fibrosis^24^. In addition, intratracheal injection, the mode of administration that is commonly used**Error! Bookmark not defined.**, is inconsistent with humans, which may lead to differences in the process of lesion formation. Furthermore, studies have shown that some therapeutic agents, although validated in BLM-induced animal models, have not been proven to be equally effective in clinical trials and even had toxic effects^25^. In consideration of these issues, we propose that, for the first time, TR might be a better approach to generate an animal model of PF.

#### 2.2. The advantages of the TR model

The advantages of the new TR model are as follows. First, the TR-containing liquid was administered by gavage, which does less damage to the mice than the BLM model, which requires anesthesia and surgery. The reduction in damage was supported by the mortality results. When the modeling step is simpler and less harmful, there are fewer factors that can interfere with the experimental results. The graph of the mortality (Figure 1C) shows that the TR model did not cause animal death during the dynamic observation after the end of modeling, while the BLM model caused an increasing number of animals to die due to the initial acute alveolitis and the subsequent rapid PF. Although the mortality rate of the BLM model gradually decreased as the condition stabilized, approximately one-third of the animals died by the experimental endpoint. Human PF rarely causes a large number of deaths at the beginning of the disease. From this perspective, this new model better models human disease than the old model. Second, the progress of fibrosis in the TR model was slow but steadily increased compared with the BLM model; thus, the TR model was more comparable to human PF^26^. Finally, both TR and BLM are popular antitumor drugs; these drugs can induce PF and can up/downregulate some indicators, which suggests that these two drugs may be similar, which requires exploration.

### 3. The proposed mechanism of TR-induced PF in mice

An obvious trend was that TR significantly decreased the activity of SOD and GSH-PX in the lung tissue of mice. After 4 weeks of modeling, the MDA content was significantly higher in the TR group than in the control group and the BLM model, indicating that TR may have induced long-lasting and severe oxidation reactions, which continually generated a large number of free radicals in the lung tissue. However, this inference contrasts with the results reported in the existing literature^27^.

The other trend was that the number of white cells in the BALF of the TR model was relatively stable during the experiment and was generally at the same level as the control group. The only type of cells that exhibited a quantitative change was neutrophils, which showed a similar trend as in the BLM model. Studies have shown that neutrophils can promote the differentiation of lung fibroblasts in culture into a myofibroblast phenotype, thereby increasing the expression of connective tissue growth factor, collagen production and fibroblast proliferation or migration^28^. Meanwhile, neutrophil elastase, a protease that is released by neutrophils, has been shown to directly cause pulmonary fibrosis after BLM-induced lung injury^29^. We speculate that TR acts on certain signaling pathways associated with neutrophils to cause high levels of elastase expression and to induce PF. It is very important to intervene with this process to verify the validity of the TR model, so we need to perform further experiments and deeper exploration in the future.

